# Speed-accuracy trade-offs in Roulette betting

**DOI:** 10.1101/2025.03.04.641449

**Authors:** Siheng Chen, S Ganga Prasath, Andrew Marantan, L Mahadevan

**Affiliations:** School of Engineering and Applied Sciences, Harvard University, Cambridge MA 02138; Department of Applied Mechanics & Biomedical Engineering, Indian Institute of Technology Madras, Chennai 600036; Department of Physics, Harvard University, Cambridge MA 02138; Department of Organismic and Evolutionary Biology, Harvard University, Cambridge 02138

## Abstract

To investigate real-time decision-making in a simplified context, we analyze the trade-offs individuals make using a game of Roulette. In this game, players observe a ball that gradually decelerates as it moves around a circular track, and the participants must predict where the ball will ultimately stop. Players are rewarded for making rapid, accurate predictions, while slower but accurate predictions, or fast yet inaccurate ones, result in lower rewards. In this speed-accuracy trade-off setup we can calculate the optimal timing for placing a bet based on the ball’s initial speed and the deceleration rate, given the capacity of an individual’s tracking abilities. We find that the participants improve their performance by accruing more reward with each trial and saturates close to the optimal performance, indicating that individuals learn the parameters of the physical model as well as a representation of the form of the reward under constrained time. Looking at the correlation between the bet time of the participant and the optimal betting time, the response delay and reward accrued, we find that the participant can be classified on a novice-expert spectrum. Our study offers ways to quantify human adaptation in competitive and dynamic environments such as sports, which may be helpful in enhancing participant performance.

## I. INTRODUCTION

Human decisions carry direct consequence for their survival and are often in situations that coerce them to make trade-offs by weighing the costs and benefits of different feasible scenarios. These decisions are made on the fly as the individual makes measurements in real-time, imposing time constraint on the process. From deciding when to hit the brakes while driving to carefully timing button presses in a rhythm game, humans make real-time decisions on a daily basis. Scientists have long suggested that human visual system plays an important role in making such decisions by resolving ambiguities in a scene by collecting and integrating sensorimotor information [1, 2] while accounting for various delays associated with proprioception, motor planning and execution [3, 4].

The mental planning to make these decisions is complex as it involves different sensory inputs each with intrinsic biases and measurement errors [5]. Recent cognitive studies suggest that the mental processes governing these decisions may be guided by simulations of the physical world akin to Newtonian dynamics [6–12] however with intrinsic uncertainty [13]. In these simulations, prior versions of the Newtonian ‘models’ are updated with new measurements/observations to improve the decisions being made. An important aspect of this decision making process is that the time-scale associated with the decision is often bounded by the duration of the dynamics of the task. Thus evaluating the costs and benefits of different actions need to be fast as demanded by the time constraints. Though there is a developing understanding of how we model the physical world based on Newtonian dynamics and the sources of noise, we are yet to understand the learning process by which we update our priors over the course of repeated trials especially when the learning has to happen over a short time-scale.

Here we look at the dynamics of decision making in a time-constrained setting through a simplified game of Roulette. In our version of Roulette, a player has to predict the stopping location of a ball slowing down on a circular track. The player’s decision is coupled to a speed-accuracy trade-off (SAT) [14–16] as their performance is associated with a reward. The reward the player gets is determined using a combination of the speed and accuracy of the bet i.e. the faster and more accurately the player predicts the stopping location of the ball, the higher is the reward; however, if the stopping location is poorly predicted or accurately predicted very late, the reward is low. The SAT paradigm provides a platform to design experiments where a player’s decision is influenced by two scenarios and is obliged to make a trade-off between them. In order to gain maximum reward the player has to make the right trade-offs between speed and accuracy as better decisions are often made after accumulating enough information about the dynamics of the slowing ball. Beyond integrating the dynamics, the player also needs to learn the features of the reward. Though SAT has been extensively used to understand the process of arriving at a decision [17], it has not been applied to a time-constrained setting where humans make decisions while simultaneously learning a representation of the physical model and the associated reward. In addition, it remains a major challenge to probe the scenarios in which an observer saturates to the ideal observer, who can optimally process and respond to speed and accuracy pressure [3, 18–20].

Through our experiments with 143 participants on the SAT task (of 35 trials), we find that the average performance of a player is limited by their capacity to measure the observables in the experiments. However, we find that the performance of each individual improves with each trial and saturates close to the optimal performance. The saturated performance of the player is determined by the intrinsic measurement limitations. In order to calculate the optimal performance for an individual, we use a Bayesian framework by coupling the posterior distribution to the dynamics of the process where the priors of the distribution are updated as the parameters of the model are learned such that it optimizes the reward function. We further find that each individual can be classified on an expert-novice spectrum based on their response delays, correlation between their betting time and the optimal betting time as well as the reward they accumulate. Overall, our results provide initial insights into the learning process when humans are forced to make trade-offs beyond the ones they already make in acquiring information through their sensory modalities.

## II. ROULETTE - AN APPARATUS TO STUDY SPEED-ACCURACY TRADE-OFF

Roulette is a classic betting game in which players bet on the characteristics (either the exact number or whether it is odd/even or the range or the color) of the final position of a rolling ball along a circular track. The more accurate a bet is (for example, betting on number 10 vs betting on a even number), the higher is the reward-ratio. Typically there are obstacles and multiple wheels on the track with the inner wheel rotating faster, making it hard to predict the ultimate position of the ball. With a completely random landing position, the casino has an edge of about 5% against the players. However, if the circular track is smooth without obstacles, and the inner wheel with the numbers are not spinning too fast, it is possible to guess the range of final position of the ball (there is usually a final time after which no more bet can be placed). Inspired by this, we use a simplified version of the Roulette where a player needs to bet on the final stopping position of a rolling ball on a circular track to probe the underlying process of perception and decision making involved in deciding the bet location.

In our version of the Roulette (shown schematically in Fig. 1C) a ball moves on a circular track with a constant drag force and the player has to predict the final position of the ball as it slows down (described in detail in sec. II C). The reward associated with the bet decays rapidly with time, forcing the player to bet quickly. Moreover the reward decreases with error: the further away the bet location is from the landing position the lower is the reward. Thus the player needs to trade off between betting fast and accurately, having enough time to watch the dynamics of the ball while not waiting for too long before the possible reward decays away - an optimal betting time exists and might be different based on each player’s perception and inference ability. Further, using the game of Roulette as our model system we design two subsidiary mini-games to identify the acumen of each player’s angle and speed measuring capability, described in the ensuing sec. II A, II B (see also Fig. 1A, Fig. 1B). Through our estimates of each player’s acuity we understand their decisions in their bets in the game of Roulette, as supported by the theoretical model (discussed in sec. II C). We performed these experiments using the online platform Prolific^TM^ with a total of 143 players. An OSF project with our initial hypothesis prior to the study is available here - [21]. Ultimately each player enrolling to participate plays a total of 3 games (see SI sec. S2 for further details, sec. S2 SS3 for the interface of the game): (*i*) Angle measuring mini-game (15 trials); Speed measuring mini-game (15 trials); (*iii*) Roulette (35 trials).

**FIG. 1.**
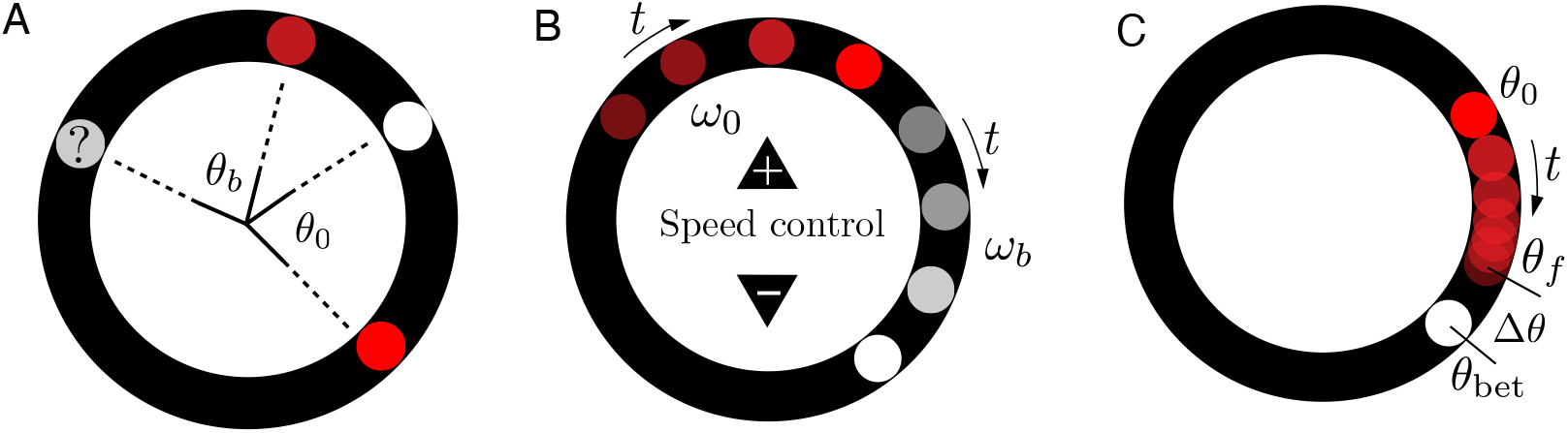
Mini-games and Roulette setup. Schematics of different games in our experiments. (A) *Angle measuring mini-game:* We first show the player a red ball and a white ball at an angle *θ*_0_ apart. We then hide this visual from the player and rotate it by a random angle. Finally, we show the picture to the player again with only the red ball. The player must then attempt to place the white ball at the same angle relative to the red ball as in the original picture. The score depends on how close the player’s angle *θ*_*b*_ was to *θ*_0_. (B) *Speed measuring mini-game:* Players first observe the red ball moving at a constant speed *ω*_0_. Then the red ball disappears and they are instructed to match the speed of the white ball *ω*_*b*_ to that of the red ball using the speed controller based on memory. (C) *Roulette game:* Players place a bet *θ*_bet_ on the final position *θ*_*f*_ of a moving red ball that starts at *θ*_0_ and experiences a constant drag force. The reward is based on both the speed of the bet and accuracy of the bet determined by Δ*θ*.

### A. Angle measuring mini-game

In the angle measuring experiments, players measure the angle spanned between a red ball (landmark ball, as in Fig. 1A) and a white ball confined to a circular track. The players are allowed to view the initial positions of both the balls for two seconds (screenshots of the game are shown in sec. S2 SS3). The two balls are then hidden from the players and rotated by a random amount. The location of the landmark ball on the track reappears afterwards, and the player is asked to place the white ball so that the angle between the two balls remains the same as before. There are 5 practice trials and 10 recorded trials for each player. We sample the initial angle, *θ*_0_ from a uniform distributed between 0 and *π*. We record the initial angle (*θ*_0_) and final angle (*θ*_*b*_) between the two balls (see Fig. 1A), as well as the response time from each player.

To quantify the angle perception error, we calculate the difference between the initial and final angles formed by the two balls, Δ*θ* = *θ*_*b*_ − *θ*_0_. Intuitively, the angle measurement error should be small when the angle is very close to 0 or *π*, since that corresponds to two balls overlapping or being on the opposite side of the circular track. Indeed we see that the betting error is close to 0 when *θ*_0_ is close to 0 or *π* (see Fig. 2A). For initial angles between 0 and *π*, players make larger errors. In particular, between *π/*2 and *π*, there is a larger variance in the betting angle, and players tend to underestimate the initial angle. If we only consider the absolute angle error, defined as |Δ*θ*| = |*θ*_*b*_ − *θ*_0_| (shown in Fig. 2B), which blind to the direction of bets we see that the error first increases then decreases when the initial angle varies from 0 to *π*.

**FIG. 2.**
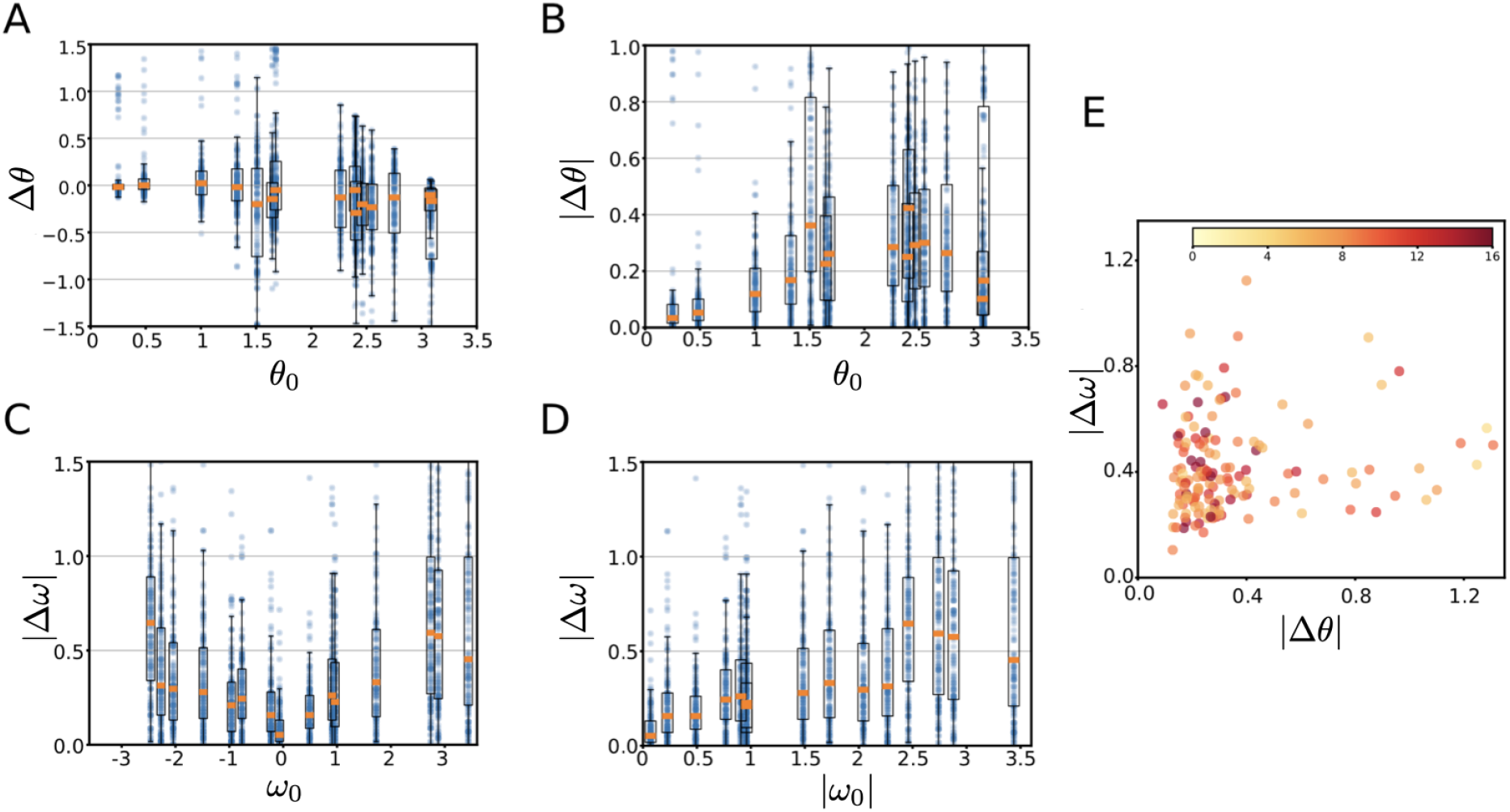
Angle and speed matching performance. (A) Distribution of angle error (Δ*θ* = *θ*_bet_− *θ*_0_) as a function of initial angle among all players. The box indicates the range between 25% and 75% quantile. (B) The absolute angle error |Δ*θ*| as a function of initial angle. (C) Absolute speed error Δ*ω* as a function of *ω*_0_. (D) Absolute speed error as a function of absolute initial speed (without directional information). (E) Speed error vs Angle error of all the players colored based on the average waiting time in the angle and speed matching games.

### B. Speed measuring mini-game

The second mini-game calibrates each player’s speed measuring acumen. The player first observes a white ball moving on a circular track with a constant speed (ref. Fig. 1B). After three seconds, the white ball is hidden for 0.5 seconds and a motionless red ball is presented along with controllers to adjust the speed of the ball (see sec. S2 SS3 for screenshots). The players are instructed to match the speed of the white ball to the speed of the red ball based on their memory using the speed controllers. There are 5 practice trials and 10 recorded trials for each player. We record the initial displayed speed of the red ball *ω*_0_ and the matched speed *ω*_*b*_ from the player, and their error defined as |Δ*ω*| = |*ω*_*b*_ −*ω*_0_|. We find that people make small errors for small initial speeds (see Fig. 2C) which increases with increases in speed, consistent with earlier observations [22]. This is more evident when |Δ*ω*| is plotted against the absolute speed, |*ω*_0_|. Beyond the increase in the magnitude of error, a commensurate increase in the variance with increase in speed is also evident. We can write *σ*_*ω*_*M* = *λω*_0_ where *σ*_*ω*_*M* is the standard deviation in the speed measurement error and *λ* is the slope captured using a curve-fit to the data for each player. This plays an important role in identifying the Roulette betting performance in the next section.

### C. Game of Roulette

After the two calibration mini-games, the players are introduced to the simplified version of the Roulette (shown in Fig. 1C). The game begins with a timer indicating when the round will start and once it begins, a red ball appears on the track from a random initial position (sampled from a uniform distribution along the track) (also ref. sec. S2 SS3). Each player gets 5 practice trials and 35 actual trials. The ball starts moving with a random initial angular position, *θ*_0_ and velocity, *ω*_0_ (sampled from a normal distribution with equal likelihood for clockwise and counterclockwise rotation). The ball itself is subject to a linear drag with a frictional coefficient of 1*/τ*_*R*_ and approaches its final position asymptotically. The evolution of the ball on this circular track with a given initial velocity *ω*_0_ has an exponentially decaying angular velocity,

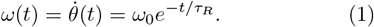

From this the position of the ball as a function of time is

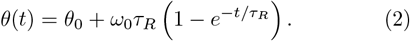

As *t* approaches infinity the final position of the ball is *θ*_*f*_ = mod (*θ*_0_ +*ω*_0_*τ*_*R*_, 2*π*) and we instruct the players to predict this final position *θ*_*f*_. Once the ball is launched, the player can use the mouse to move a circular cursor around the track to indicate *θ*_bet_ as the angle they predict to be the final position *θ*_*f*_ of the ball. After the player clicks the mouse, we record the betting position *θ*_bet_ along with the time the player took to make the bet, *t*_*b*_. The reward, *R*(*t*_*b*_) for the bet placed by the player is a function of both the accuracy and the time taken to make the bet; the faster and more accurate the player makes the prediction the higher is the reward. The functional form of *R*(*t*) is defined by

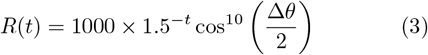

where Δ*θ* = (*θ*_*f*_ − *θ*_bet_) is the prediction error and *θ*_bet_ is the player’s prediction of *θ*_*f*_. The even function cosine accounts for the periodicity of the *θ*(*t*) and also that the reward is symmetric about Δ*θ* → − Δ*θ*. The exponent 10 is added such that the function is peaked at *θ*_*f*_ and penalizes bets far away (see SI Fig. S1 for the shape of the reward). The player needs to find their own trade-off based on the perception and inference capabilities so that the reward is maximized.

#### Calculating optimal betting time

The optimal betting time, *t*_opt_ maximizes the rewards *R*(*t*) given the intrinsic capabilities of the individual. The dynamics of a ball relaxing with a time-scale *τ*_*R*_ from Eqs. 1, 2 can be written in the form **s**(*t*) = *U* (*t*; *τ*_*R*_)**s**_0_ where **s**(*t*) = (*θ*(*t*), *ω*(*t*)) and the initial condition **s**_0_ = (*θ*_0_, *ω*_0_) with

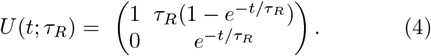

We are interested in the final position of the ball *θ*_*f*_ and it can be written in terms of the initial condition as *θ*_*f*_ = **a**^*T*^ **s**_0_ such that **a**^*T*^ = (1, *τ*_*R*_). Since *θ*_*f*_ is a linear function of the initial condition **s**_0_ (which is chosen as a Gaussian with mean ***µ***), it follows that *θ*_*f*_ is also Gaussian with mean

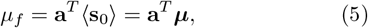

where ⟨⟩denotes the average. We see that the estimate of *θ*_*f*_ through *µ*_*f*_ depends on the player’s measurements through ***µ*** (reff SI Eq. S15). Now in order to derive an expression for the player’s estimation of **s**_0_, we first need to capture the properties of the measurements made by the player. Let the vector **s**_*M*_ (*t*) represent the measurements we make of the state **s**(*t*) from time 0 to time *t*. For our purposes here, we assume **s**_*M*_ (*t*) is a Gaussian signal centered about the actual state, **s**(*t*) with correlation function *C*_*M*_ (*τ, τ* ^′^) (see SI sec. S1 SS1 for a detailed derivation),

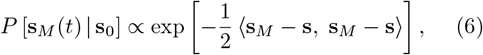

where we have used the notation,

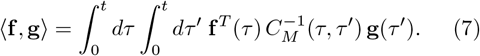

We argue the functional form of *C*_*M*_ (*τ, τ* ^′^) below, and further use Bayes’ theorem, which is the key to calculating *P* (**s**_0_| **s**_*M*_ (*t*)), the probability distribution of the initial state **s**_0_ given measurements **s**_*M*_ (*t*). Using Bayes’ theorem, we can write *P* (**s**_0_ | **s**_*M*_ (*t*)) in terms of *P* [**s**_*M*_ (*t*) | **s**_0_] (in Eq. 6) and our prior distribution for **s**_0_ as,

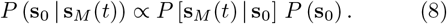

We know that *P* (**s**_0_) is a multivariate Gaussian which we use to sample the initial angle, *θ*_0_ and angular velocity, *ω*_0_ with mean ***µ***_0_ and variance Σ_0_ given by

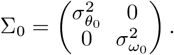

Thus we need a functional form for *C*_*M*_ (*τ, τ* ^′^) to close the system and estimate the distribution *P* (**s**_0_ | **s**_*M*_ (*t*)). We assume that the correlation function is proportional to the measurement error capabilities (captured by the variance in angle measurement error, 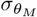 and speed measurement error, *σ*_*ω*_*M*) and the time period of observation *τ*_*M*_. We then get

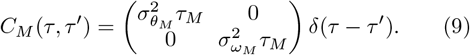

This functional form is motivated by the insight that the correlation is proportional to the duration of the observation, *τ*_*M*_ and the measurement error variances,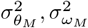.

With this assumption, we can now calculate the mean final angle, *µ*_*f*_ and the variance *σ*_*f*_ through *P* (**s**_0_ | **s**_*M*_ (*t*)) in Eq. 8 as *µ*_*f*_ = **a**^*T*^ ⟨**s**_0_⟩. After some algebra (detailed in SI sec. S1 SS2) the variance of the final angle,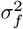, when the player has no prior information about the initial speed 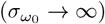 is,

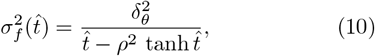

where we have introduced three dimensionless parameters (see SI Eq. S22 for details): the rescaled time 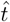, the rescaled angular measurement error 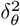 and the relative speed measurement error *ρ*^2^ defined as,

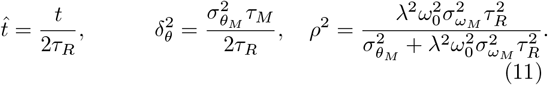

Here 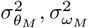 are the variance of the measured angle and speed signals, *τ*_*M*_ is the time provided for observation in the speed-measuring mini-game, while *λ* is the coefficient from a curve-fit to speed vs speed error data in Fig. 2D and has a dimension of rad^−1^. In the expression for relative speed measurement error, *ρ*^2^ we have introduced our observation in sec. II B that the absolute prediction error in angular speed is proportional to the initial speed i.e. 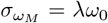 (ref. Fig. 2C, D). It is this linear relationship that brings about the dependence of the initial speed on the speed measurement error.

The probability distribution of the final position of the red ball given the measurements *θ*_*M*_ (*t*), *ω*_*M*_ (*t*), *P* (*θ*_*f*_ | *θ*_*M*_ (*t*), *ω*_*M*_ (*t*), *τ*_*R*_) contains everything the player knows about *θ*_*f*_ at time *t*. It however does not *a priori* tell us *how* to place their bet. Moreover, the player not only has to decide where to place the bet but also *when* to place it. This comes by maximizing the expected value of the reward function, given by (see SI sec. S1 SS1 for further details)

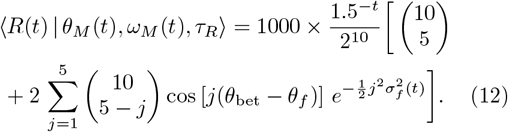

First we consider how the player should choose *θ*_bet_ given what they have measured up to time *t*. If the player knows the reward function in Eq. 3, they can place the bet to maximize their expected reward. Evidently the expression for 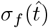 depends both on the speed measuring acumen, brought through the dependence on the initial speed, as well as the measurements made till time *t*. We see that the expected reward is largest when *θ*_bet_ = *µ*_*f*_ and thus the player should choose *θ*_bet_ at any given time *t* to match their expected value of *θ*_*f*_ at that time. If they do so, their expected reward becomes

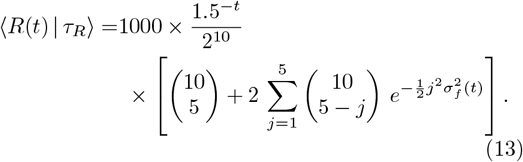

Since 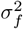 in Eq. S24 depends on the measurement of the initial speed *ω*_0_, the expected reward is a non-monotonic function of the measurement (see SI Fig. S1). Unfortunately, it is difficult to get an analytic expression for *t*_opt_ but we can compute it numerically by finding the time corresponding to the maximum in the expected reward, ⟨*R*(*t*) | *τ*_*R*_⟩:

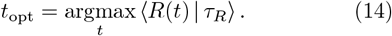

#### Performance enhancement in betting

With the optimal betting time, *t*_opt_ in Eq. 14 calculated using a numerical procedure, we can investigate how each player’s performance changes over the course different trials. We see in Fig. 3D that after a few practice rounds the players make better trade-off between speed and accuracy by betting at a time closer to their optimal betting time. By defining an optimality index *β* as the ratio between the reward the players get at a particular trial and the reward at the optimal betting time,

**FIG. 3.**
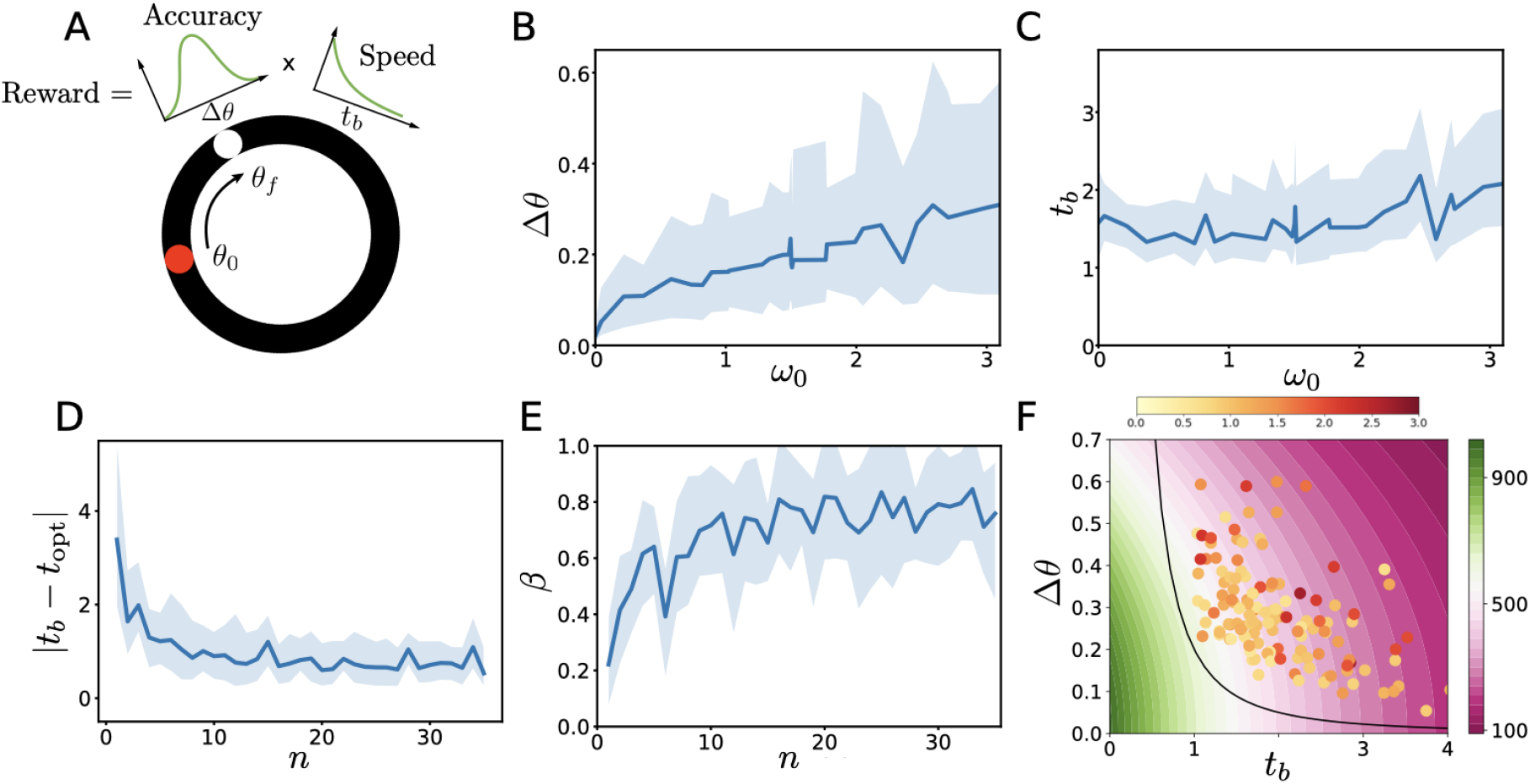
Performance in roulette game. (A) Schematic of the reward, which is a function of the difference angle error (Δ*θ*) and the betting time (*t*_*b*_) (see Eq. 3). (B, C) Betting error Δ*θ* and betting time *t*_*b*_ as a function of the initial speed of the red ball (*ω*_0_), showing the classical linear increase in error with angular speed. The betting time *t*_*b*_ also exhibits a weak linear increase with the speed. (D) Error in betting time, |*t*_*b*_ − *t*_opt_| defined as the difference between betting time, *t*_bet_ and optimal betting time, *t*_opt_ = argmax_*t*_ ⟨*R*(*t*) |*τ*_*R*_ ⟩ (using Eq. 13) as a function of trial *n*. (E) Performance index, *β* defined ratio of reward obtained by the player to optimal reward (which is the reward at the optimal betting time, *t*_opt_) as a function of trial *n* for all players. We see a clear improvement in player performance attributable to learning of the parameters of the game. (F) Betting error, Δ*θ* vs betting time, *t*_*b*_ for all players with circle color indicating their speed measuring acumen from mini-game 2. We see that larger the angle error in mini-game 1, larger is the betting error in Roulette. The solid line corresponds to Δ*θ*^2^*t*_*b*_ = 0.2 where Δ*θ* = ⟨*θ*_*f*_ − *θ*_bet_⟩. The contour is the magnitude of reward for the particular betting error, Δ*θ* and betting time, *t*_*b*_.

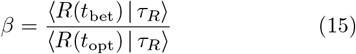

we see in Fig. 3E that *β* approaches 1 after 15 trials. Moreover it also indicates that the players learn a representation of the only independent parameter in the model i.e. the frictional coefficient 1*/τ*_*R*_ accurately. We can visualize the average performance of each player individually in the phase-space defined by betting error, Δ*θ* vs betting time, *t*_*b*_ in Fig. 3F. Note that the region closer to the origin in this phase space corresponds to a larger reward. We estimate that the error and betting time of all the players is bounded by the curve ⟨*θ*_*f*_ − *θ*_bet_⟩^2^*t*_bet_ = 0.2. Moreover we find that the speed acumen (captured through 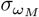 in speed measuring minigame in sec. II B) seems to matter more in determining a player’s performance than their angle measuring acumen, which is seen through the color of circles in Fig. 3F, where a brighter color corresponds to large errors from the speed measuring mini-game in sec. II B.

Average performance *β* helps us understand the general trend in learning and inference in the physical process, but the individual response can provide further insight into what distinguishes between a skilled player from an unskilled player. Plotting the betting time, *t*_*b*_ and optimal betting time, *t*_opt_ for each individual player (see Fig. 4A left panel), we see that the difference between them is larger in the beginning than in the end. Some players tend to follow the optimal betting time well (Player 16), while some do not (Player 20) or even have a betting time inversely correlated with the optimal betting time (Player 19). In addition, some players tend to wait longer to bet (Player 20 and 13), while some do not (Players 16 and 19). These can all be captured by computing the slope of the fit between the betting time, *t*_bet_ against the the player’s optimal betting time, *t*_opt_ (ref. Fig. 4A right panel). By doing a simple linear regression between *t*_bet_ and *t*_opt_, we can calculate the R-squared *r*^2^, slope *k*, and intercept *b*, which indicates the player’s tendency to follow the optimal betting time, the delay factor (for longer *t*_opt_ how much larger *t*_bet_ will be assuming zero reaction time), and the player’s reaction time. Intuitively if a player follows the optimal betting time, has a small delay factor, and has a short reaction time, this player should perform well. When we combine these three factors as *r*^2^*/*(*kb*), we see in Fig. 4B that it has a clear positive correlation with the reward *R* capturing the average performance (correlation of *r*^2^ = 0.41). This quantity can be used to discern a player’s performance quality and can help classify them either as a novice or an expert for large or small values of *r*^2^*/*(*kb*).

**FIG. 4.**
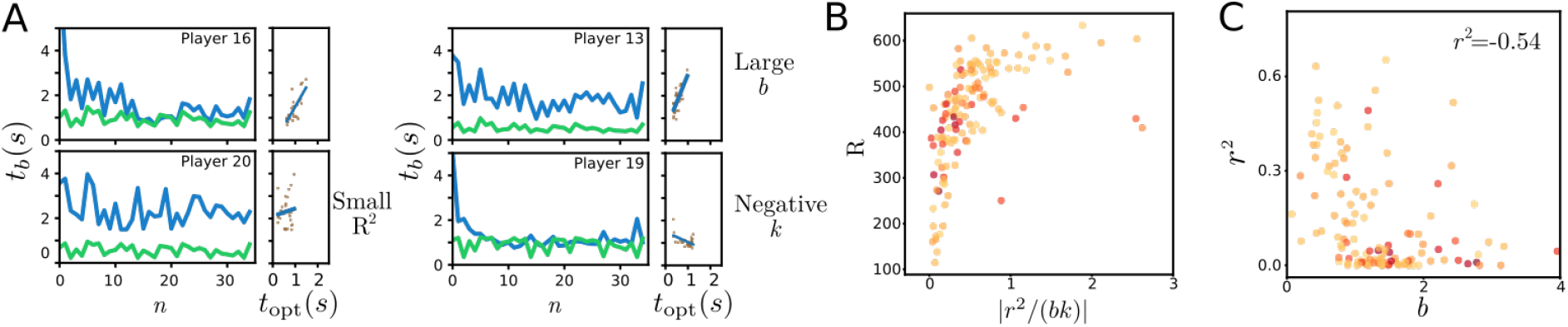
Player performance. (A) Example trajectories of players showing the betting time, *t*_*b*_ (green curve) following (left panels) and not following (right panels) the optimal betting time (blue curve) in the roulette game. For each player, the plot on the right sub-panel shows the correlation between the betting time, *t*_*b*_ and the optimal betting time, *t*_opt_. (B) Roulette reward, *R* vs |*r*^2^*/*(*bk*) | for all players, where *r*^2^, *k* and *b* are the R-squared, slope, and intercept of the linear fit between *t*_*b*_ and *t*_opt_. (C) Scatter plot of *r*^2^ and *b* of all players. Color shows the age of the players.

The comparison between betting time, *t*_*b*_ and optimal betting time, *t*_opt_ also sheds light on how humans make adjustments to maximize their reward. It has been shown that humans can adjust the amount of mental simulations to optimally balance speed and accuracy [23]. In the roulette game, we also see a similar adjustment: the players who have a longer response time (larger intercept in the regression of *t*_bet_ vs *t*_opt_) do not follow the optimal betting time very well (smaller *r*^2^) (shown in Fig. 4C). This can be understood as follows: people with a longer response time compensate for their slowness by making a bet faster. Furthermore, slower learners (smaller *k*) tend to have a similar waiting time for both difficult and easy trials, which is more evidence of the compensation effect in how humans trade-off speed and accuracy.

## III. CONCLUSION

In this article, through the simple task of predicting the final position of a ball slowing down from its initial speed on a circular track, we studied the decision-making process in humans. By coupling the predictions that the players make to a reward and asking them to maximize this reward, we investigate how they learn the parameters associated with the dynamics of the ball. Over the course of our experiments, we found that the players learned the parameters of the model to increase their reward the limit of which is set by their capacity to accurately track the ball.

The relationship of a player’s betting times (*t*_bet_) to their optimal betting times (*t*_opt_) captures the *total delay* in the mental simulation and how the delay changes with the difficulty of the task through the slope. It also provides an estimate of the response time when no simulation is required through the offset *b* (for example, when *ω*_0_ = 0). It is perhaps these subtle trade-offs under time-constrained settings that distinguish experts from novices in the context of competitive activities such as sports and our approach provides a template for making quantitative estimates.

## Supplemental Information for

## S1. THEORETICAL MODEL

### SS1. Inference of linear dynamical systems

What do we know about the properties of a linear dynamical system after measuring it over some period of time? Let us define some notation to make this more precise. Let the vector **s**(*t*) denote the state of the system at time *t*. Since we are assuming a linear dynamical system, we can describe the evolution of the state with time as

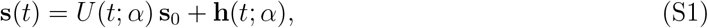

where **s**_0_ denotes the initial state, *U* (*t*; *α*) is the time-evolution matrix, **h**(*t*; *α*) is a state-independent vector related to external forcing terms and *α* represents all the parameters upon which the state evolution depends. Finally, let the vector **s**_*M*_ (*t*) represent the measurements we make of the state from time 0 to time *t*. For our purposes here, we assume **s**_*M*_ (*t*) is a Gaussian signal centered about the actual state with correlation function *C*_*M*_ (*t, t*^*′*^),

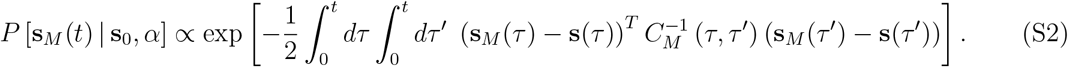

Note that the inverse correlation function above represents an operator inverse. Note also that we use square brackets to denote a *functional* distribution. It will be useful at this point to introduce the following inner product notation,

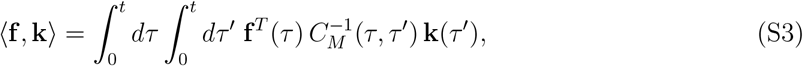

so that we can write Eq. S2 more compactly as

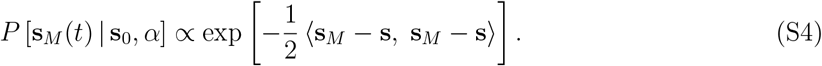

Now we can address our first question: we can express our state of knowledge about the system parameters (**s**_0_ and *α*) given the measurements **s**_*M*_ (*t*) via the probability distribution *P* (**s**_0_, *α* | **s**_*M*_ (*t*)).

The key to calculating *P* (**s**_0_, *α* | **s**_*M*_ (*t*)) is Bayes’ theorem, which lets us write it in terms of *P* [**s**_*M*_ (*t*) | **s**_0_, *α*] (Eq. S2) and our prior distribution for **s**_0_ and *α, P* (**s**_0_, *α*),

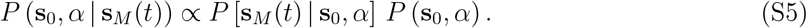

Here we make two assumptions about our prior. First, we assume **s**_0_ and *α* are independent, so that

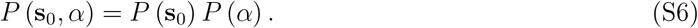

Second, we take *P* (**s**_0_) to be multivariate Gaussian,

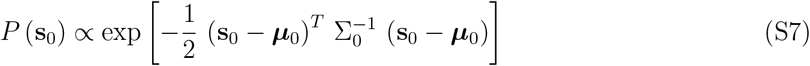

Given the Gaussian form of both Eq. S2 and Eq. S7, it turns out that *P* (**s**_0_, *α* | **s**_*M*_ (*t*)) is also Gaussian in **s**_0_. To see this, we first expand the quadratic form in the prior,

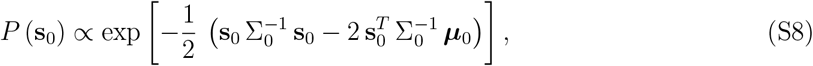

where we absorbed the term with no **s**_0_ or *α* dependence into the proportionality sign. Then we expand the inner product in the measurement term and substitute in Eq. S1 to get

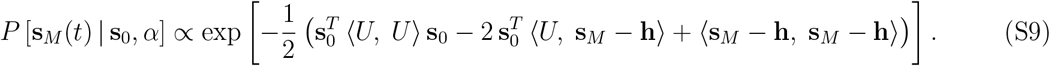

Note that we are able to take **s**_0_ outside of the inner products as it is constant in time. Upon substituting the above expressions into Eq. S6 and collecting terms in **s**_0_, we obtain

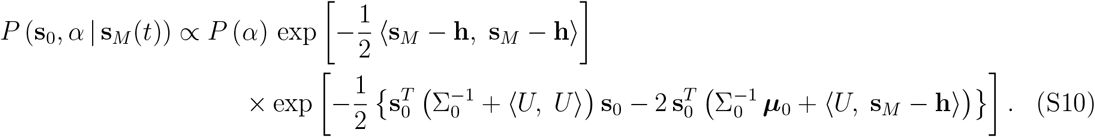

Here it useful to introduce some new notation. We let Σ^*−*1^(*α*) represent the inverse covariance matrix corresponding to the measurement term,

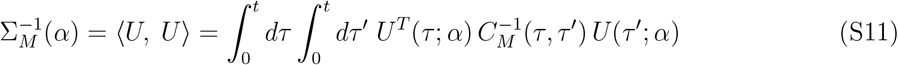

and we let ***µ***_*M*_ (*α*) represent the most likely state vector according to the measurement part,

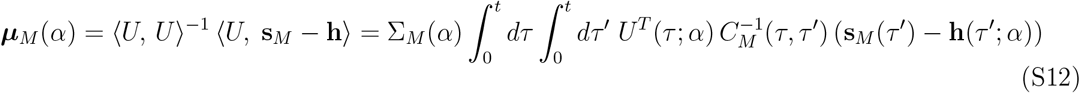

Using this new notation, Eq. S10 becomes

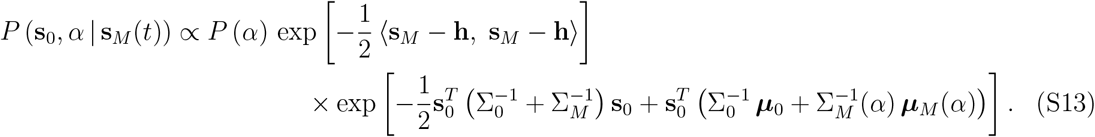

Then, in preparation for completing the square in **s**_0_, it makes sense to define

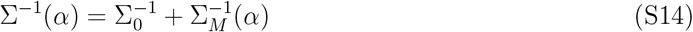

and

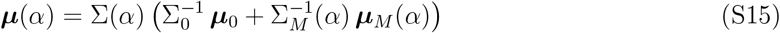

so that Eq. S13 simplifies to

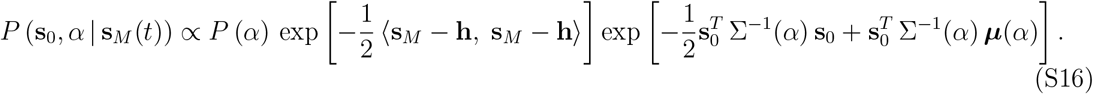

Completing the square then yields our final result,

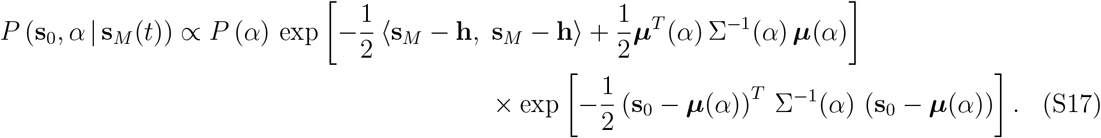

While our posterior is multivariate Gaussian in **s**_0_, it is not the case for the other system parameters *α*. Fortunately the fact that **s**_0_ appears as multivariate Gaussian makes it easy to calculate the marginal distribution for *α*,

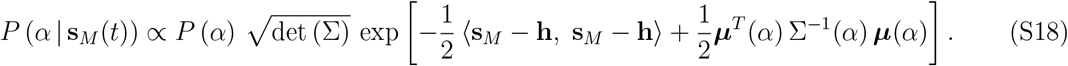

### SS2. Game of Roulette

In our simplified game of Roulette, we ask the player to predict where a decelerating ball will finally stop along the track given by *θ*_*f*_. The earlier the player makes a bet, the higher the potential reward. The point of this game is to investigate how players trade off speed (placing bets earlier) and accuracy (waiting longer to make more accurate bets). The game begins by flashing a “Get ready!” message to the player while a visible timer indicates when the round will start. We then launch the ball on the track from a random initial position, *θ*_0_ and with a random angular velocity, *ω*_0_ drawn from a Gaussian distribution with a randomly chosen sign. The ball itself is subject to linear drag and only approaches its final position asymptotically. The game of Roulette then is given by the dynamics of the ball relaxing with a time-scale *τ*_*R*_. The state vector defined by the angular position and velocity along the track, **s**(*t*) = (*θ*(*t*), *ω*(*t*)), is simply:

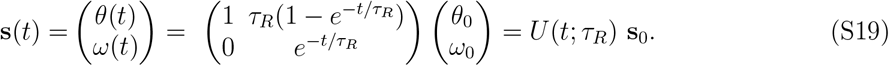

The variable of interest in this game is the final position of the ball, given by *θ*_*f*_ which is the angle that the ball subtends when *t*→ ∞. From the above equation we have *θ*_*f*_ = lim_*t→∞*_ *θ*(*t*) = *θ*_0_ +*ω*_0_*τ*_*R*_. This can be written in term of the initial condition simply as *θ*_*f*_ = **a**^*T*^ **s**_0_ such that **a**^*T*^ = (1, *τ*_*R*_). The mean value of this final angle can be evaluated using Eq. S15.

Once the ball is launched, the player is free to use the mouse to move a circular cursor around the track to indicate what they predict to be the final position of the ball. Once the player clicks the mouse, we record the bet, *θ*_bet_ along with the time the player took to make the bet, *t*_*b*_. Finally, we show the player their reward, defined by

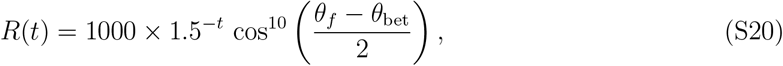

where *θ*_*f*_ is the final resting angle of the ball and *θ*_bet_ is the player’s prediction of *θ*_*f*_. Note that we have chosen the reward to fall by a factor of 2/3 for every second that passes. We expect this to be more meaningful for players to understand than a more general exponential decay with time.

### SS3. Inference

Given the dynamics of the ball in Roulette described by the state vector **s**(*t*) = (*θ*(*t*), *ω*(*t*)), we expect the player is able to learn a representation of the only parameter in the system which is the decay rate *τ*_*R*_ as well as the behavior of the reward *R*(*t*). Further we expect that the player can make accurate measurements of the angle and angular velocity of the ball along the track. This assumption means that the player’s distribution **s**_*M*_ (*t*) of **s**(*t*) at time *t* given their measurements of the position and the velocity to be multivariate Gaussian.

However in the roulette game, what really matters is the player’s state of knowledge about the final angle *θ*_*f*_. We can obtain the expression for *θ*_*f*_ by taking the limit of *θ*(*t*) as *t* → ∞, yielding *θ*_*f*_ = *θ*_0_ + *ω*_0_ *τ*_*R*_. Letting **a**^*T*^ = (1, *τ*_*R*_), we can write this as *θ*_*f*_ = **a**^*T*^ **s**_0_, and since *θ*_*f*_ is a linear function of the Gaussian **s**_0_, it follows that *θ*_*f*_ is also Gaussian with mean

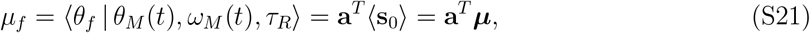

which depends on the player’s measurements through ***µ*** (from Eq. S15), and variance

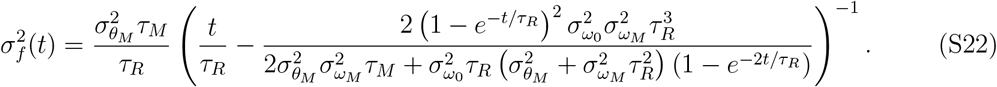

which does not depend at all on what the player measures (from Eq. S14). In order to obtain this expression, we have assumed *C*_*M*_ (*τ, τ* ^*′*^) of a particular form

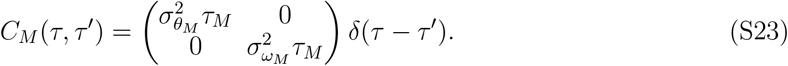

The variance becomes much more manageable when the player has no prior information about the initial speed (i.e. in the limit as 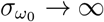), reducing to

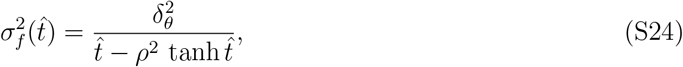

where we have introduced three important dimensionless parameters: the rescaled time 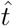, the rescaled angular measurement error 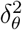 and the relative speed measurement error *ρ*^2^. The expressions for these three are

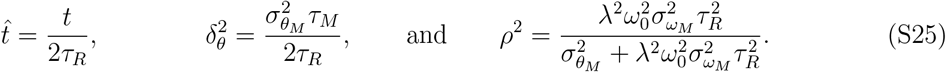

These parameters also make it easier to write down the form of the mean of *θ*_*f*_,

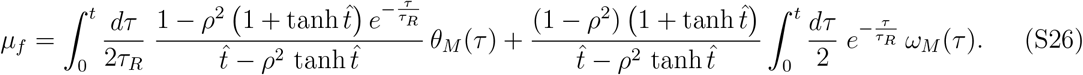

### SS4. Optimal betting time

While *P* (*θ*_*f*_ | *θ*_*M*_ (*t*), *ω*_*M*_ (*t*), *τ*_*R*_) contains everything the player knows about *θ*_*f*_ at time *t*, it does not *a priori* tell them *how* to place their bet. Moreover, the player not only has to decide what bet to place, but also *when* to place it. Here we go through this process, starting by asking what the player should choose as *θ*_bet_ if they were to place a bet at time *t*. In the course of doing this, we will show that, so long as the player knows *τ*_*R*_, their expected reward at time *t* will not depend on what they have measured, and so they can determine at what time to place their bet based on when the expected reward is largest.

First we consider how the player should choose *θ*_bet_ given what they have measured up to time *t*. If the player knows the reward function in Eq. 3, they can place their bet to maximize their expected reward:

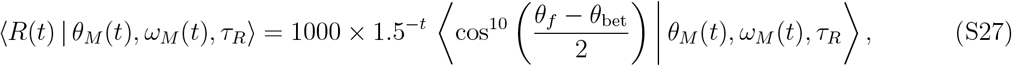

We can simplify the above expression using the complex identity

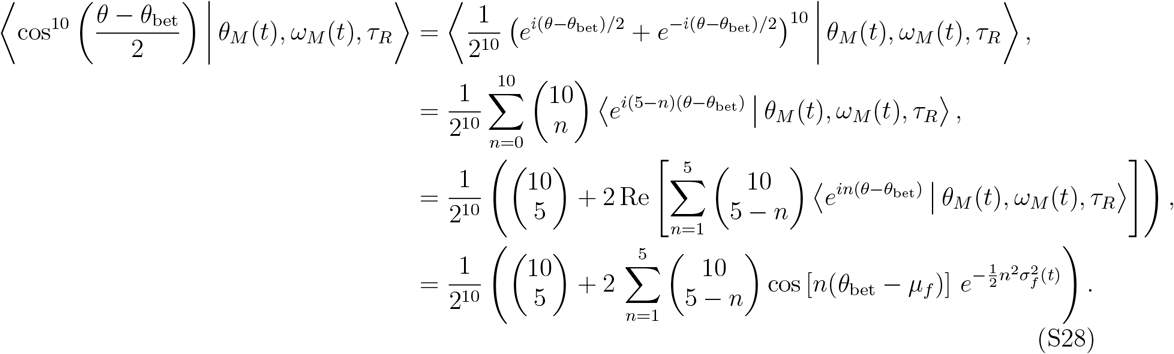

Evidently the expression in Eq. S28 and hence the expected reward is largest when *θ*_bet_ = *µ*_*f*_. Thus the player should choose *θ*_bet_ at any given time *t* to match their expected value of *θ*_*f*_ at that time. If they do so, their expected reward no longer depends on the measurements and becomes. We thus get

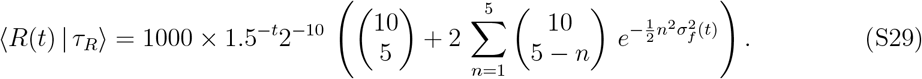

We can compute the time corresponding to the maximum in the expected reward, ⟨*R*(*t*) | *τ*_*R*_⟩ by numerically solving for *t*_opt_ such that it satisfies

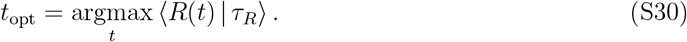

### SS5. Visualizing betting strategies

To visualize the position of the optimal betting time, we set the parameters to two sets of values: (1) 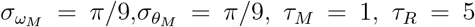 (2) 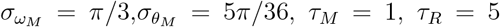. And the resulting expected reward as a function of betting time are shown in Fig. S1A and S1B respectively. We can see that with certain combination of parameters, which correspond to certain angle and speed perception capability, the optimal betting time is nonzero (as shown in Fig. S1A), suggesting that observing the evolution of the ball helps improving the performance. For some other combinations of parameters, it is in fact more preferable to place a bet directly in the beginning (see Fig. S1B), since the time cost of visual perception outweigh the gain in the betting accuracy.

**FIG. S1.**
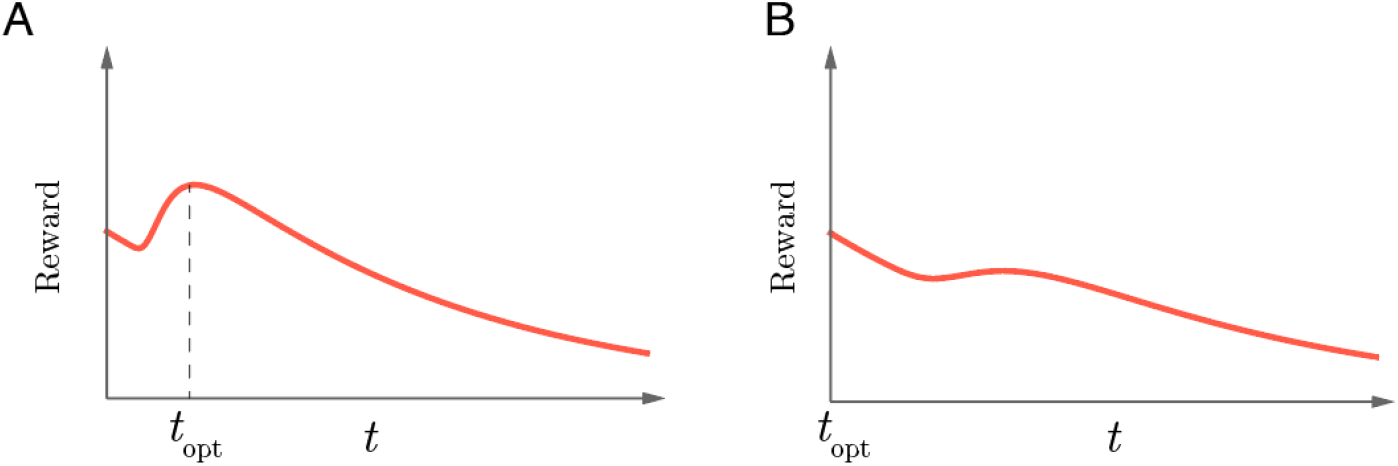
(A) When the measurement error in angle and speed perception is small, the optimal betting time is non-zero. Waiting for a few seconds to gain information results in a better bet (higher reward). (B) If the measurement error is too large, it is better to bet immediately.

This difference in optimal betting time can indeed be observed in different players. In Fig. S2, the betting time (blue) and the optimal betting time (green) for the 35 roulette game trials (including the first 5 practice trials) for 27 players are shown. For each user, the linear regression between the betting time and the optimal betting time is also shown on the right. For player 41 and 81, the optimal betting time suggests that for some initial speed of the ball, the optimal strategy is to bet without observing the dynamics of the ball. Player 81 seemed to have followed this suggestion in some trials.

## S2. EXPERIMENTAL DETAILS

### SS1. Parameters in experiments

- Angle error (15 trials): Initial angle: Uniform distribution between 0-2*π*. Rotation angle: Uniform distribution between 0-2*π*. Presentation time: Present: 2 sec, Disappear: 0.5 sec.
- Speed error (15 trials): Normal distribution with *µ* = 5*π/*9, *σ* = 5*π/*18 of which 50% is clockwise and the rest anti-clockwise. Presentation time: Present: 3 sec, Disappear: 0.5 sec.
- Roulette (35 trials): We drew the initial angle from a uniform distribution between 0-2*π*. For initial speed we used a normal distribution with *µ* = 5*π/*9, *σ* = 5*π/*18 of which 50% is clockwise and rest is anti-clockwise. The drag time decay *τ*_*R*_ was fixed at 1.5*s* for the first 25 trials and then drawn from a Gaussian with *µ* = 1.5*s* and *σ* = 0.5*s* for the final 10 trials.

### SS2. Demographic information

First, the age and performance of players are negatively correlated (Fig. S4A, *R*^2^ = − 0.14). This could result from poorer capability in trading off between speed and accuracy, or age-related reductions in dexterity or vision. Second, the number of years of education does not seem to have any effect on the performance (Fig. S4B, *R*^2^ = − 0.001). Third, the male players seem to perform better than female players (Fig. S4C, *p* = 0.003). Finally, the handedness does not affect the performance significantly (Fig. S2D, *p* = 0.39).

**FIG. S2.**
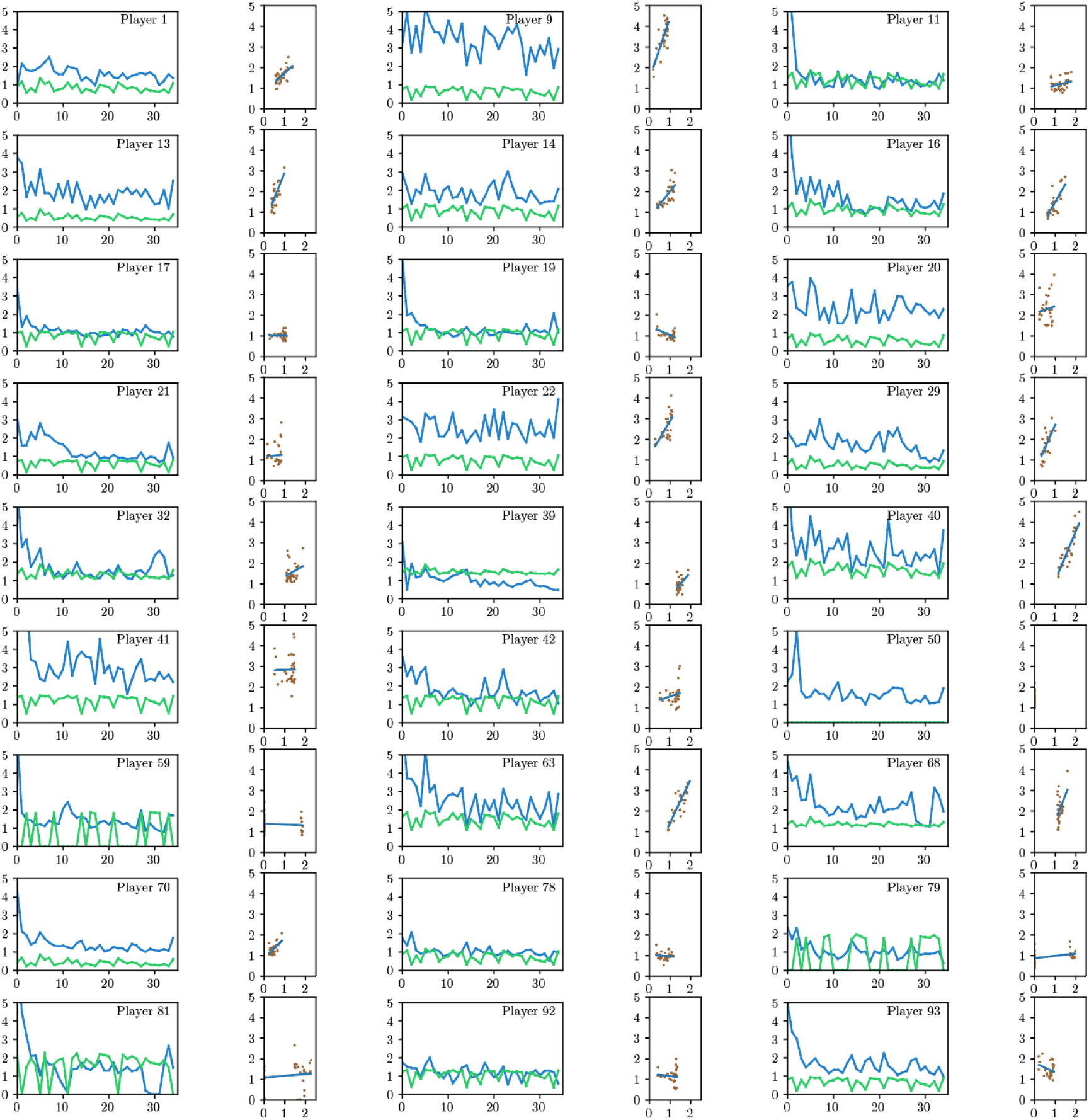
The betting time (blue) and optimal betting time (green) as a function of the number of trials, for 27 selected players. For each player, the betting time vs optimal betting time is shown on the right (scatter plot). Many players follow the optimal betting time. For some people, the change in their betting time is opposite to the change in the optimal betting time. Some user has optimal betting time being zero (poor measurement in the previous games), and one person tried to follow this zero optimal betting time (Player 81, betting immediately).

### SS3. Interface design of the game

Fig. S5 shows the screenshots of the game interface for the angle matching game and the speed matching game. Fig. S6 shows the screenshots of the roulette game.

**FIG. S3.**
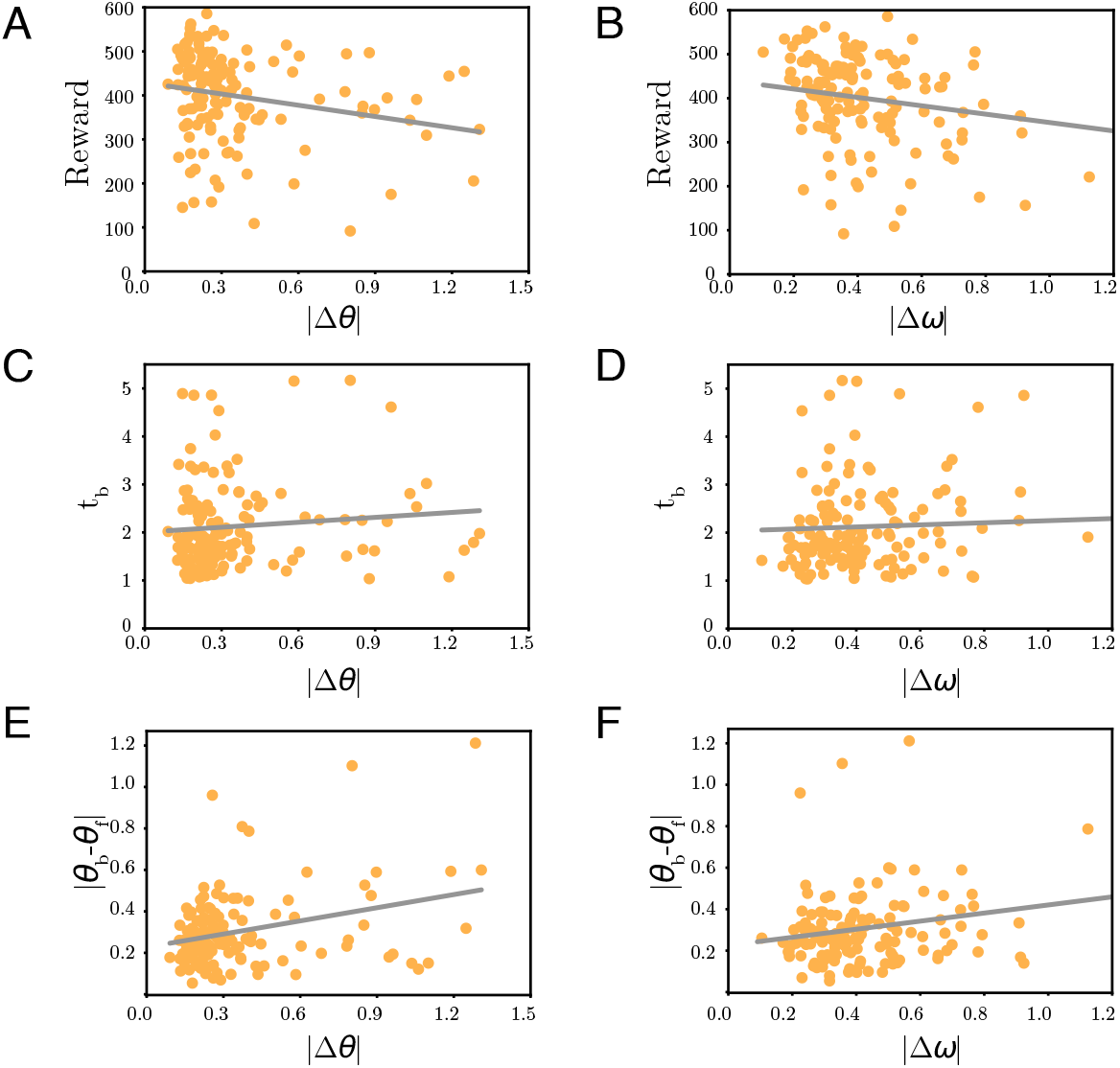
Relationship between the average reward in the roulette game and the average angle error (A) or speed error (B). Each point represents one participant in the experiment. (C) and (D) show the relationship between the betting time in the roulette game and the angle error / speed error respectively. (E) and (F) show the angle error in the roulette game w.r.t. angle / speed error respectively.

**FIG. S4.**
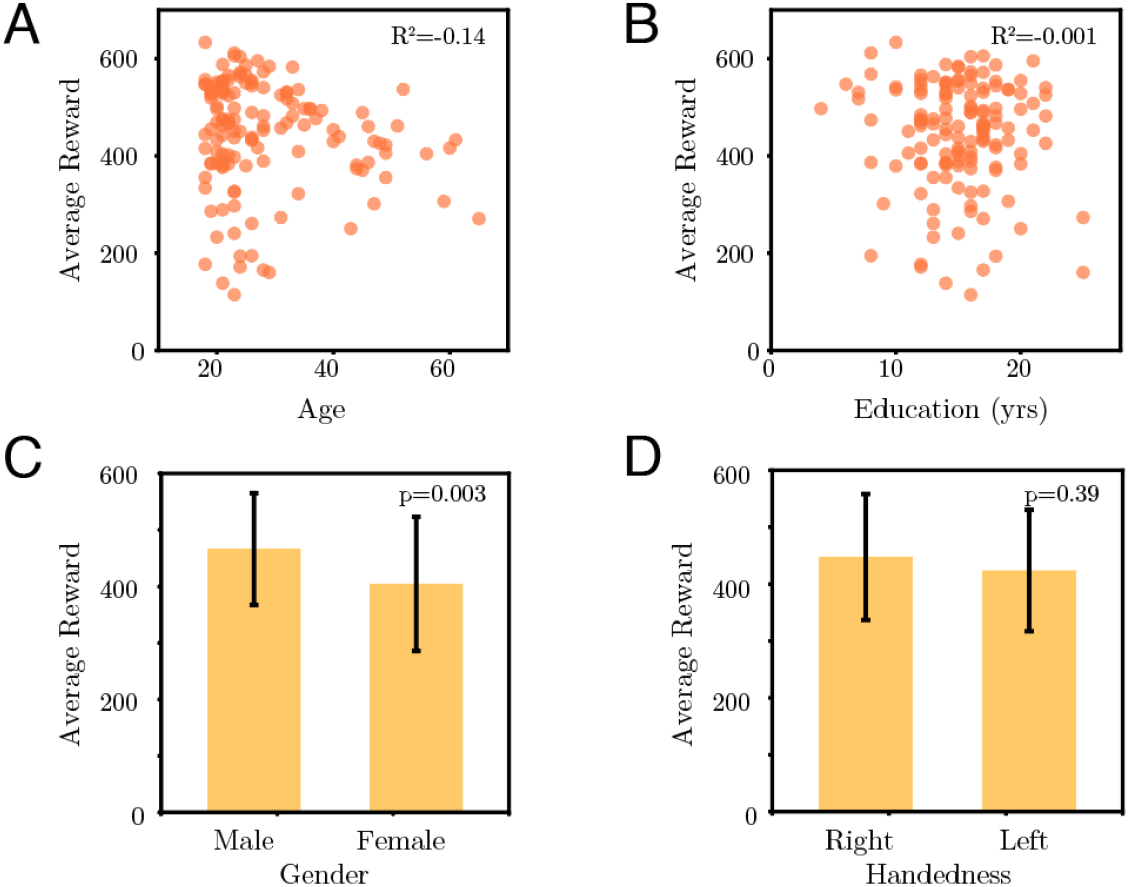
Effect of age, education, gender, and handedness on the roulette performance.

**FIG. S5.**
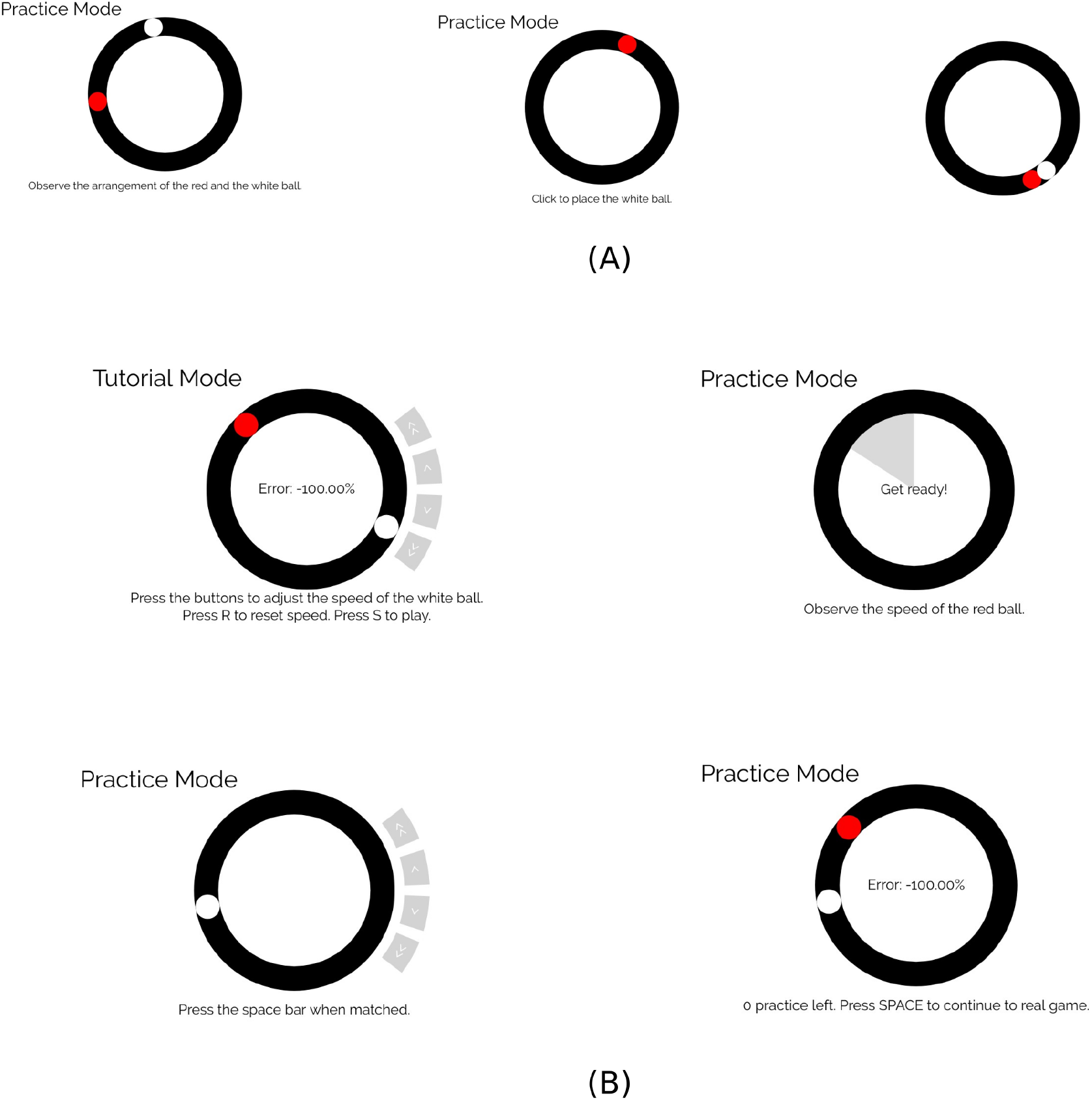
Screenshots of (A) The angle mini-game and (B) the speed mini-game.

**FIG. S6.**
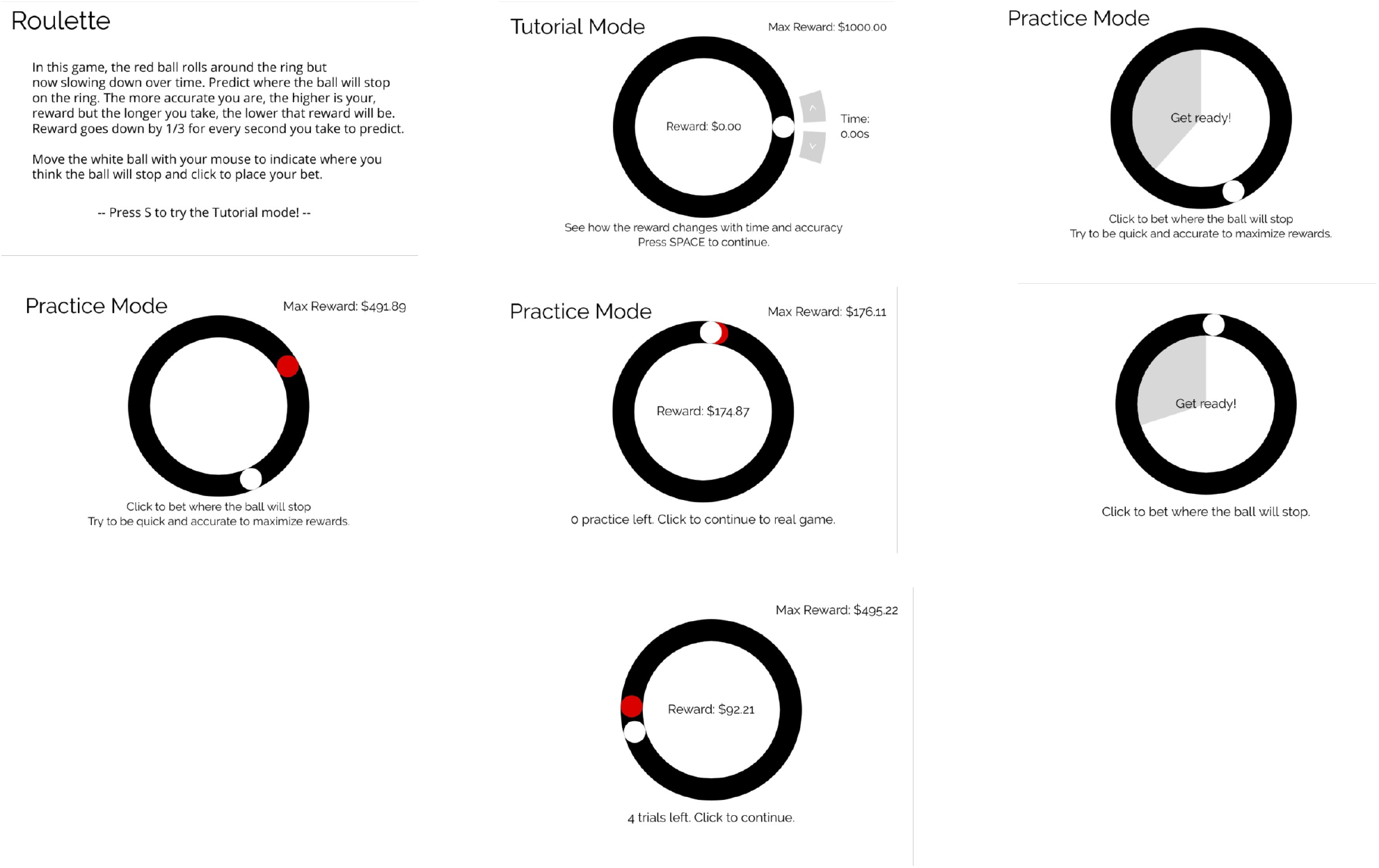
Screenshots of the Roulette game.

